# L-type Voltage-gated Calcium Channel Modulators Inhibit Glutamate-induced Morphology in Astrocytoma Cells

**DOI:** 10.1101/2019.12.15.875799

**Authors:** Mitra Sadat Tabatabaee, Frederic Menard

## Abstract

The excitatory neurotransmitter glutamate evokes physiological responses within the astrocytic network that lead to fine morphological dynamics. However, the mechanism by which astrocytes couple glutamate sensing with cellular calcium rise remains unclear. Employing natural properties of U118-MG astrocytoma cells, we tested a possible connection between L-type voltage-gated calcium channels (Ca_v_) and glutamate receptors. Using live confocal imaging and pharmacological inhibitors, the extension of U118-MG processes upon glutamate exposure are shown to depend mainly on extracellular calcium entry via L-type Ca_v_’s. Inhibitors of the Ca_v_ α1 protein, decreased astrocytic filopodia extension; while, gabapentinoids, ligands of the Ca_v_’s α2δ auxiliary subunit blocked all process growth. This study suggests that α2δ is the main contributor to Ca_v_’s role in glutamate-dependent filopodiagenesis. It opens new avenues of research on the role of α2δ in neuron-astrocyte glutamate signaling and neurochemical signaling at tripartite synapses.

## INTRODUCTION

Astrocytes have historically been described as nonexcitable cells that do not contribute actively to information processing in the brain, in contrast to neurons who communicate actively through electrical or chemical synapses. However, this simplistic view has evolved over the past two decades as more detailed studies on astrocytes have emerged (Araque et al. 1999; Cornell-Bell, Thomas, and Caffrey 1992; Rose et al. 2018; Bazargani and Attwell 2016). For instance, astrocytes are now known to respond to glutamate among other stimuli (Cornell-Bell, Prem, and Smith 1990; Rose et al. 2018; Cornell-Bell, Thomas, and Caffrey 1992). This astrocytic glutamate sensitivity led to the tripartite synapse model, where astrocytes are active participants in a functional neural connection (Araque et al. 1999). However, the molecular mechanisms underlying this signaling are still not fully understood (Rose et al. 2018).

Glutamate is the main excitatory neurotransmitter released at neuronal synapses (Rose et al. 2018). In astrocytes, glutamate triggers an intracellular calcium rise, followed by a rearrangement of the actin cytoskeleton, which results in movement, formation, and protrusion of filopodia (Cornell-Bell, Thomas, and Caffrey 1992). These filopodia in close proximity to synapses *in vivo* are termed peripheral astrocytic processes (PAPs) (Molotkov et al. 2013; Santello, Toni, and Volterra 2019). They are highly motile structures that seek to contact synapses (Haber, Zhou, and Murai 2006). Several factors have been demonstrated to influence PAPs motility and filopodia formation: neuronal activity, synapse-specificity, mGluR5 presence, Ca^2+^ concentration, and IP3 concentration (Bernardinelli et al. 2014; Perez-Alvarez et al. 2014; Heller and Rusakov 2015).

Filopodia formation in astrocytes upon glutamate stimulation has been proposed to occur via two main pathways based on both ionotropic glutamate receptors (iGluR) and metabotropic glutamate receptors (mGluR) (Cornell-Bell, Thomas, and Caffrey 1992; Rose et al. 2018; Heller and Rusakov 2015; Matyash and Kettenmann 2010). The most abundant astrocytic iGluRs are the calcium-permeable AMPA receptors, and they have been proposed to control glutamate-dependent filopodiagenesis (Bernardinelli, Muller, and Nikonenko 2014; Heller and Rusakov 2015). Alternatively, mGluR5 has also been proposed to be the main receptor responsible for this phenomenon (Bernardinelli et al. 2014; Heller and Rusakov 2015; Panatier and Robitaille 2016). In both cases, glutamate sensing was shown to correlate with an intracellular calcium rise in astrocytes. With Ca^2+^-permeable AMPA receptors, glutamate causes a calcium influx from the extracellular milieu (Iino et al. 2001). With mGluR5, exposure to glutamate releases Ca^2+^ from internal stores through 1,4,5-inositol trisphosphate (IP3) signaling (Panatier and Robitaille 2016). In both pathways, the calcium rise leads to actin cytoskeletal rearrangement combined to filopodiagenesis, processes extension and motility (Heller and Rusakov 2015).

**Figure 1.**
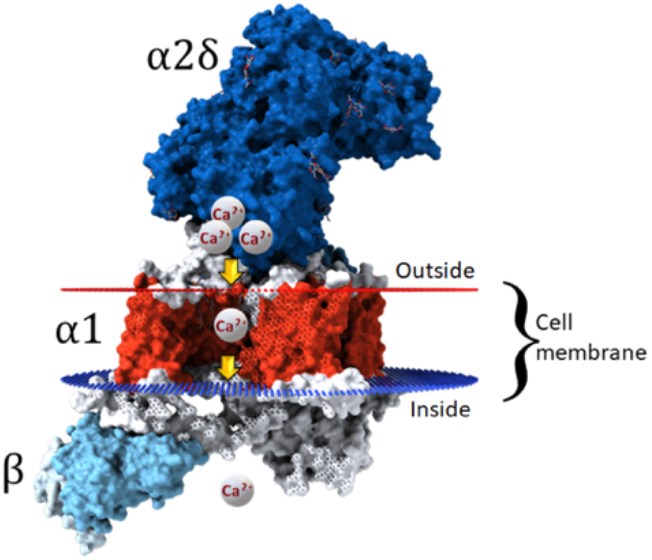
Assembly of L-type Ca_V_ channels. The α1 subunit (red) is composed of 24 transmembrane helices that form the selective Ca^2+^ pore. The intracellular β subunit (light blue) is responsible for the channels trafficking to the membrane. The extracellular α2δ subunit (dark blue) is the modulator of α1 subunit conductance and controls the localization of the channel on the cell membrane (modified from PDB structure 5GJV).

During the course of an unrelated study, we observed that glutamate-induced filopodiagenesis was affected by voltage-gated Ca^2+^ channel (Cav) modulators in the astrocytoma cell line U118-MG. The *human protein atlas* shows that U118-MG express only two types of glutamate receptors: mGluR8 and an isoform of AMPA receptor possessing iGluR2 subunits (GRIA2). Neither of these can directly link glutamate sensing with the calcium rise that mediates the filopodia extension. Consequently, we investigated whether a connection exists between glutamate sensing and *high voltage-gated calcium channels* (L-type Ca_v_’s) in U118-MG cells.

Cav’s are classified according to their electrophysiological and pharmacological properties into high Ca_v_’s (L-, N-, P-, Q-, and R-type) and low Cav’s (T-type) (Latour et al. 2003; Zamponi et al. 2015). Each channel couples a change in membrane potential to a Ca^2+^ influx, where Ca^2+^ ions act as secondary messengers in events such as contraction, secretion, and gene expression (Woolley et al. 1990). Structurally, they consist of three main subunits that associate in a 1:1:1 stoichiometry: a pore-forming α1 subunit, an external α2δ subunit, and an intercellular β subunit (Figure1) (Zamponi et al. 2015; Thul et al. 2017). The β subunit ensures the Cav channel localizes to the membrane, while the α1 and α2δ subunits regulate ion conductance. The transmembrane α1 forms the calcium channel pore and contains the voltage sensor. The extracellular α2δ subunit modulates properties of α1 such as its activation and conductance rate (Woolley et al. 1990; Striessnig and Koschak 2008; Song et al. 2015).

Astrocytes are not electrically excitable cells, yet they express L-type Ca_v_’s (Latour et al. 2003). The exact function of these voltage-dependent channels in astrocytes is still unknown, but their contribution in pathological conditions of the brain has been extensively demonstrated, with Cav1.2 in particular (Cheli et al. 2016; Wang et al. 2015; Westenbroek et al. 1998; Anekonda and Quinn 2011; Shaw and Colecraft 2013; Daschil et al. 2013). The only L-type Ca_v_ shown to be expressed in U118-MG cells is Cav1.2 (Thul et al. 2017).

We originally hypothesized that Cav might participate to filopodiagenesis in astrocytes when the cells are subjected to glutamate stimulation by amplifying intracellular Ca^2+^ concentrations after initial activation of iGuR and/or mGluR. iGluRs and phospholipase Clinked mGluRs (group II and III) are known to cause an intracellular Ca^2+^ rise upon glutamate binding that can lead to the gating of Cav’s (Verkhratsky and Kirchhoff 2007). Adenylate cyclase-linked mGluRs (group I) can also trigger the gating of Ca_v_’s by activating the channel through phosphorylation (Hove-Madsen et al. 1996). In both cases, gating of Cavs can provide enough calcium to support the cytoskeletal reorganization that leads to a morphological change. Therefore, *we expected* that interrupting calcium conductance of Ca_v_ would inhibit or decrease the morphological response to glutamate in astrocytes.

In this study, we initially showed that the calcium required for filopodiagenesis in response to glutamate is supplied from the extracellular milieu, which suggests an active role of membrane ion channels. Next, to test our hypothesis on the role of Cav’s, we employed antagonists for the pore-forming α1subunit of L-type calcium channels and ligands selective for its α2δ subunit. We were surprised to find that the effect of α2δ ligands was stronger than the effect of α1 antagonists. In fact, gabapentinoids targeting α2δ completely inhibited the morphological response of U118-MG cells to glutamate, while α1 antagonists only partially decreased the cellular response. Herein, we present evidence suggesting that the α2δ auxiliary subunit is the main contributor to Cav role in the glutamate-dependent filopodiagenesis of astrocytes.

## MATERIALS AND METHODS

### Cell culture

Human astrocytoma U118-MG cell line (HTB-15, ATCC) was gifted by Dr. Andis Klegeris, UBC Okanagan. U118-MG cells were cultured in Dulbecco’s modified essential medium (DMEM, Gibco 11995-065) supplemented with 10% v/v heat-inactivated fetal bovine serum (HyClone 12483020) 1% v/v penicillin 10,000 U/ml and streptomycin 10,000 μg/ml (Gibco 15140-163). Cells were incubated in a humidified atmosphere containing 5% CO_2_ at 37 °C, and typically passaged at 80-95% confluence using a pre-warmed 0.25% trypsin-EDTA solution for a maximum of 5 minutes (Gibco SH30236.02).

For experiments, cells from a 10 cm culture dish at ~80% confluence were resuspended with a pre-warmed 0.25% trypsin-EDTA solution and transferred to 35 mm glass-bottom culture dishes (Mutsunami D1130H) in phenol red-free, high glucose DMEM supplemented with 25 mM HEPES (Gibco 21063-029). Enough cells were transferred to obtain a ~50% confluent culture dish after adhesion. Cells were incubated for less than the cells’ doubling time, typically ~33 h to minimize imaging variation between samples (see statistical analysis section) (Westermark 1973). This period allows cells to adopt a flattened shape that helps define the filopodia for measurements. Identical incubation times were respected to ensure cells were at the same growing stage in all treatments.

Prior to imaging, the medium was saturated with CO_2_ bubbling delivered via a needle attached to a 0.22 μm syringe filter. CO_2_ maintain U118-MG viability on the microscope (Aumann et al. 2017).

### Transfection

U118-MG astrocytoma cells were transfected with 1200 ng/μl of the F-actin marker mCherry-LifeAct-7 plasmid using a calcium phosphate precipitation protocol (Kingston, Chen, and Rose 2003). mCherry-Lifeact-7 was a gift from Michael Davidson (Addgene plasmid # 54491)

### Cells treatment

On the microscope stage, U118-MG cells in phenol red-free, high glucose DMEM supplemented with 25 mM HEPES (Gibco 21063-029) were stimulated with glutamate (Alfa Aesar A15031-30). Briefly, 10.0 μL of a 20X glutamate stock solutions as carefully delivered to the medium (2.0 mL in a 35 mm dish) at the outside edge of the dish for a final concentration of 100 μM. The stock solution of L-glutamate was prepared in deionized millipore water and filter-sterilized through a 0.22 μm syringe filter (Cornell-Bell, Prem, and Smith 1990; Aumann et al. 2017).

For Ca^2+^-blocking experiments, 1.00 μL of a 200X stock solution of Cav inhibitor was delivered to the glassbottom culture dishes (Mutsunami D11130H) for a final concentration of each reagent as shown in Table 1. After each addition, cells were incubated at 37 °C under a humidified 5% CO_2_ atmosphere for the time indicated in Table 1 (4 minutes to 3 hours).

Stock solutions of Cav inhibitors were prepared in DMSO (vehicle) before each experiment. Solutions were filter-sterilized using a 0.22 μm syringe filter. To avoid negative effect of DMSO on the cells, stock solutions were highly concentrated (200X), thereby maintaining the final amount of DMSO below 1% v/v in the media. The exception were gabapentinoids, which were prepared directly in deionized millipore water vehicle, also as a 200X stock solution for consistency.

For calcium chelation experiments, either intracellular or extracellular, the same protocol as for Cav inhibitors was used with times and concentrations listed in Table 1. Confocal microscopy and live imaging

Imaging of live cells was conducted on an Olympus FV1000 fluorescence confocal microscope equipped with a Plan-ApoN 60x/i.4 oil-immersion objective. Fluorescence of mCherry-Lifeact-7 was observed at 559 nm excitation using a He–Ne laser (Olympus filter set Cy3.5).

Changes in cell morphology were recorded at 60X magnification. To measure filopodia extension, an image of a single cell was acquired every 20 second for a total of number of 60 frames.

### Quantification of filopodia extension

The length of at least twenty filopodia per cell was measured manually using FIJI image analysis software. Only filopodia whose arbor could be unambiguously visualized throughout the entire imaging period were selected (Zatkova et al. 2018). A filopodium’s length was measured from the edge of the cell membrane to its apical end; each manual measurement was repeated three times on the same image. The change in length was calculated by subtracting the length of a process prior to stimulation (frame 10; three minutes) from the length of the same filopodium after 20 minutes (frame 60). Glutamate was delivered to the medium four minutes after the beginning of imaging (i.e., frame 12). The reported “change in length” for one cell represents the average of 20 measured filopodia per cell.

### Statistical analysis

Each value represents the averaged changes measured from at least 5 cells, each from a separate culture dish (n ≥ 5); the results are presented as the mean ± S.E.M. Significant differences among groups were determined using one-way or two-way analysis of variance (ANOVA) and *post hoc* analysis using Prism 7.04 from GraphPad. A *p* value < 0.05 was considered significant. Statistically non-significant data is not starred on plots. To minimize technical variations within our samples for ANOVA analysis, we employed a *completely randomized block design* (Krzywinski and Altman 2014). Briefly: cells of a dish at 80-95% confluency were trypsinized and simultaneously re-plated in several 35 mm imaging dishes and incubated in a humidified atmosphere containing 5% CO_2_ at 37 °C for less than 33 hours to avoid variation between samples.

## RESULTS

### U118-MG cells extend their processes in exposure to glutamate

When astrocytoma cells were imaged in their medium without stimulation, their filopodia length changed by 0.28 ± 0.24 μm over the course of 16 minutes. In contrast, when cells were subjected to 100 μM glutamate (4 min. after beginning of imaging), their processes grew by 4.82 ± 0.95 μM (Fig. 1B). This 17-fold increase in length was statistically significant (p > 0.0001). A rapid process extension was observable within the first two to five minutes after glutamate exposure (not shown), but the final length filopodia were measured 16 minutes poststimulation to allow cells to stabilize.

### The glutamate-induced process extension in U118-MG cells depends on extracellular calcium

Chelation of extracellular free Ca^2+^ ions with EGTA in the medium completely abolished filopodia extension upon exposure to 100 μM glutamate (*p* < 0.0001, Fig. 1B). In presence of EGTA, the change in length is statistically indistinguishable from the baseline cell response. However, when the intracellular free Ca^2+^ was sequestered with BAPTA-AM, the cell’s response to glutamate stimulation was significantly reduced, but not completely inhibited: filopodia extension was about 35% of that observed with control cells (*p* < 0.001, two-way ANOVA post hoc Sidak’s multiple comparison tests).

#### Blocking the pore-forming subunit of L-type Cav’s does not block the morphological response of astrocytoma cells to glutamate

Figure 2 displays the effect of subjecting U118-MG cells to drugs known to inhibit Ca^2+^ ion influx by interacting with the α1 subunit of L-type Ca_v_’s. Upon glutamate stimulation, the processes extension decreased significantly (*p* < 0.0001), but was still observed to be 30-50% of the negative control with unblocked cells. More specifically, when cells were pre-treated with drugs for 30 minutes prior to glutamate stimulation, the average increase in processes length was: 2.36 ± 0.77 μm with 10 μM nifedipine, 1.82 ± 0.60 μm with 10 μM verapamil, and 1.70 ± 0.79 μm with 10 μM ditiazem. Ca_v_ α1-targeting inhibitors decreased filopodia protrusion by 50-65% compared to the positive controls, i.e., untreated cells (4.82 ± 0.95 μm) or vehicle treatment (4.30 ± 0.92 μm).

**Figure 2.**
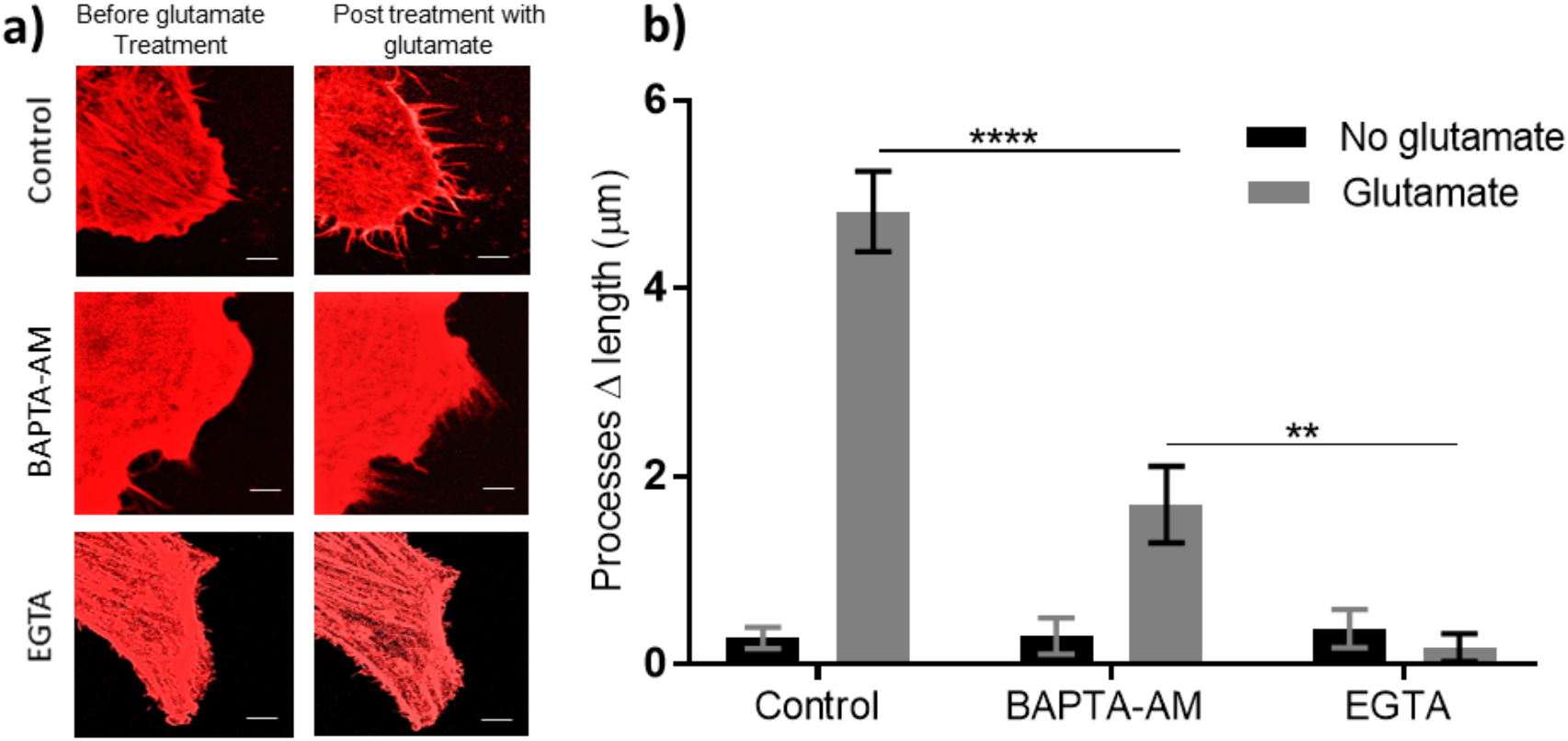
Extracellular calcium is essential for glutamate-induced filopodia extension in model astrocytes. (**a**) Representative images of U118-MG astrocytoma cells upon exposure to 100 μM glutamate for 20 minutes when treated with and without intra-or extracellular calcium chelators. Confocal fluorescence micrographs of cells transfected with mCherry-LifeAct7 at 60X magnification; scale bar is 5 μm. (**b**) Cells extend their processes significantly in response to 100 μM glutamate. Chelating intracellular Ca^2+^ (10 μM BAPTA-AM) or extracellular Ca^2+^ (50 μM EGTA) decreases the length of processes extension induced by 100 μM glutamate. Chelating the intracellular calcium with BAPTA-AM still allowed for significant process extension, yet it is significantly shorter than that of the control cells with glutamate. In contrast, chelating the extracellular calcium with EGTA inhibited filopodia growth. Change in length (Δ) was calculated by subtracting the length of a filopodium immediately prior to stimulation from that of the same filopodium after 20 minutes. Results are presented as mean ± SEM. The change in length is the average data of 5 cells, each from a separate culture dish; measurements of at least 20 filopodia were averaged for each cell (n = 5). All comparisons use a *post hoc* Tukey’s multiple comparison (two-way ANOVA, F _*2,24*_ glutamate treatment = 34.98, F _*2,24*_ Ca^2+^ chelators = 71.38, for both factors p < 0.0001), **p= 0.002, ****p < 0.0001.

#### Antagonists of L-type Cav’s α2δ subunit block the morphological response of astrocytoma cells to glutamate

Processes extension and filopodiagenesis were completely inhibited when U118-MG cells were pretreated with gabapentinoids drugs for 30 minutes, prior to glutamate exposure (Fig. 3). When 10 μM solutions of either gabapentin or pregabalin in water were used, no filopodia extension was observed upon glutamate stimulation (*p* > 0.8, one-way ANOVA Dunnett’s multiple comparison post hoc test). The changes in processes length were statistically indistinguishable from the background growth without glutamate stimulation.

**Figure 3.**
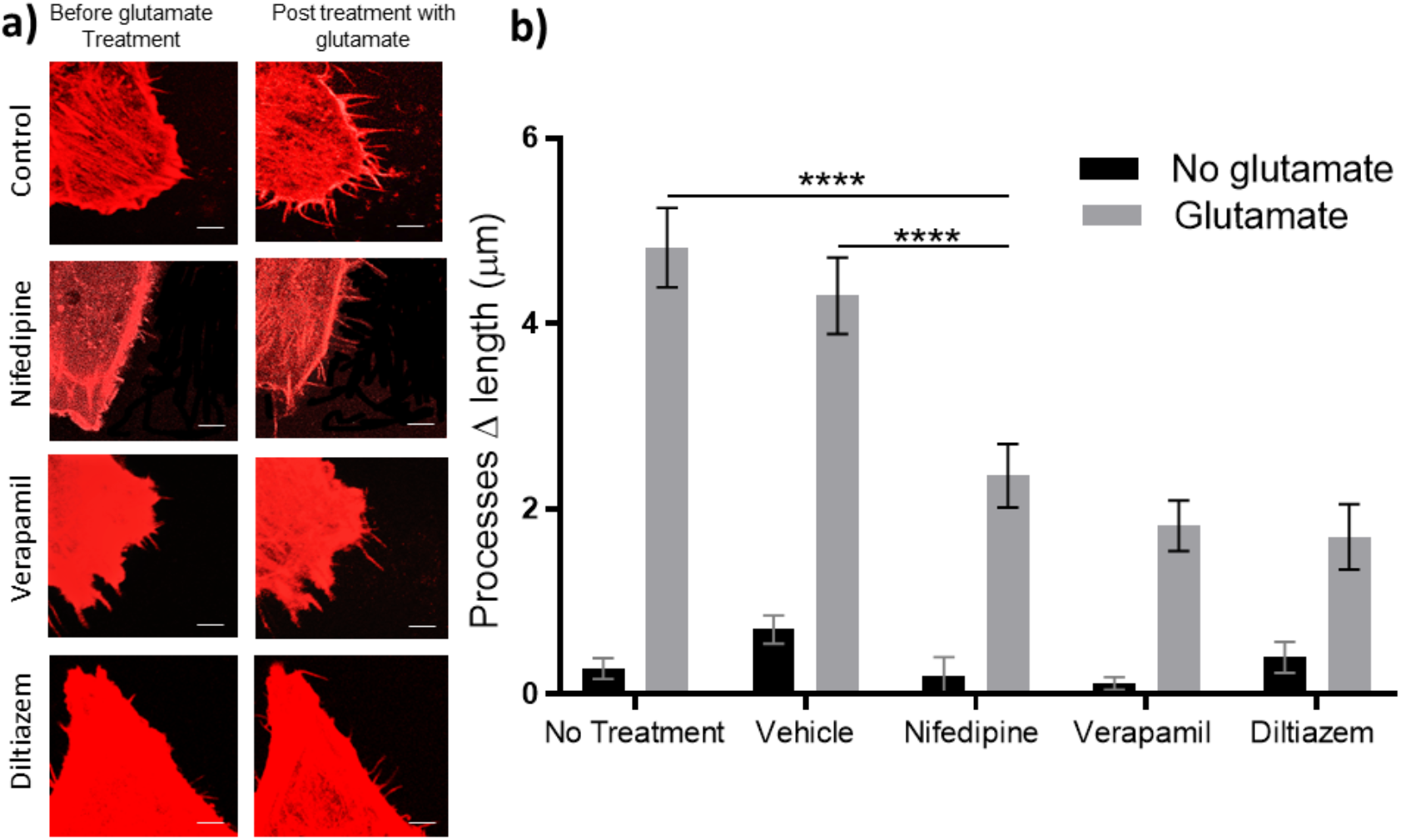
Blocking the pore-forming subunit of L-type Ca_V_’s significantly decreases processes extension induced by glutamate in U118 cells. (**a**) Representative images of U118-MG astrocytoma cells upon exposure to 100 μM glutamate after being treated with L-type CaV blockers (10 μM) for 30 minutes prior to stimulation. Confocal fluorescence micrographs of cells transfected with mCherry-LifeAct7 at 60X magnification; scale bar is 5 μm. (**b**) Blocking the pore-forming subunit of L-type calcium channels significantly decreases the processes extension upon exposure to glutamate. All treatments with glutamate caused a significant increase in processes length compared to unstimulated cells. Processes extension was significantly decreased when cells were treated with L-type Ca_V_ blockers prior to glutamate exposure. Vehicle solution was DMSO only. Change in length (Δ) was calculated by subtracting the length of a filopodium immediately prior to stimulation from that of the same filopodium after 20 minutes. Results are presented as mean ± SEM. The change in length is the average data of 5 cells, each from a separate culture dish; measurements of at least 20 filopodia were averaged for each cell (n = 5). Data were analyzed by two-way ANOVA (F _*4,40*_ glutamate treatment = 16, F _*1,40*_ Ca^2+^ blockers = 227.3, for both factors p < 0.0001). All comparisons done by *post hoc* Tukey’s multiple comparison.

**Figure 4.**
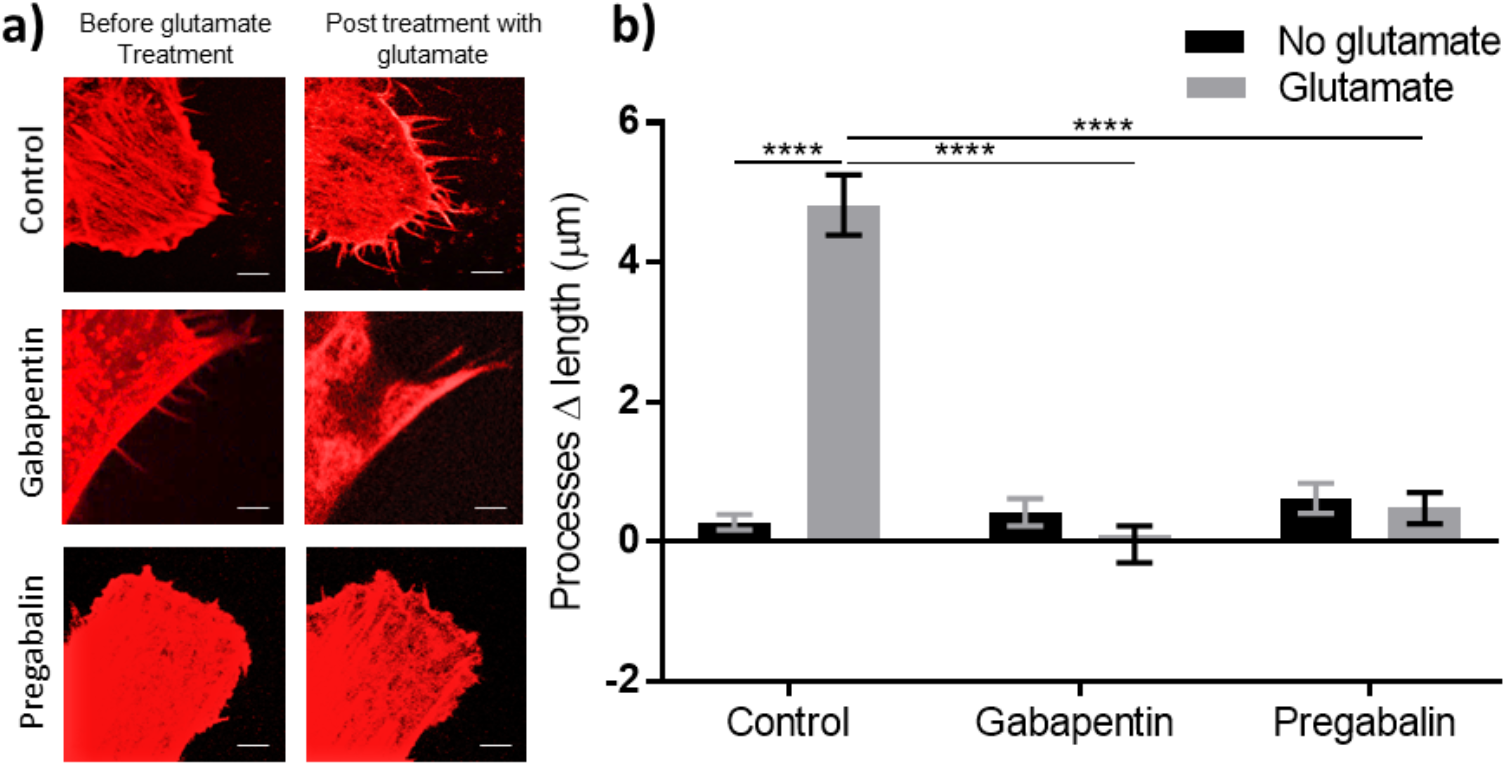
The α2δ subunit of L-type Ca_V_ controls filopodia extension when astrocytoma cells are exposed to glutamate. (**a**) Representative images of U118-MG cells exposed to 100 μM glutamate after being treated (or not) with 10 μM α2δ subunit inhibitors for 30 minutes prior to glutamate stimulation. Confocal fluorescence micrographs of cells transfected with mCherry-LifeAct7 at 60X magnification; scale bar is 5 μm. (**b**) Pharmacologically blocking the α2δ subunit of L-type Cav’s with gabapentinoids completely inhibited filopodia growth or extension in response to 100 *μ*M glutamate. Change in length (Δ) was calculated by subtracting the length of a filopodium immediately prior to stimulation from that of the same filopodium after 20 minutes. Results are presented as mean ± SEM. The change in length is the average data of 5 cells, each from a separate culture dish; Measurements of at least 20 filopodia were averaged for each cell (n = 5). Data were analyzed by two-way ANOVA (F _*124*_ glutamate treatment = 38.75, F _*224*_ treatments = 48.72, for both factors p < 0.0001); data comparisons used a *post hoc* Dunnet multiple comparison (p < 0.0001).

### DISCUSSION

In this study, L-type voltage-gated calcium channels (Cav) are shown to play a significant role in the glutamate-induced filopodiagenesis of astrocyte model cells U118-MG. Of the three protein subtypes constituting L-type Cav, the α2δ subunit appears essential to control filopodia extension, while the transmembrane channel α1 subunit may only be partially involved. These results are consistent with what has been observed in primary astrocytes (Cornell-Bell, Thomas, and Caffrey 1992; Cornell-Bell, Prem, and Smith 1990; Araque et al. 1999; Bernardinelli et al. 2014; Aumann et al. 2017). U118-MG is a well-established human astrocyte model cells, derived from malignant gliomas by Ponten in the late 1960’s along with U138-MG. The transcriptome of U118-MG and U138-MG has been characterized; they have identical variable number tandem repeat (VNTR) and similar short tandem repeat (STR) patterns (Uhlen et al. 2010).

In our study, sequestering the extracellular calcium during glutamate stimulation prevented the protrusion of cell processes; while, sequestering the intracellular calcium still elicited about a third of the response (Fig. 1). Although it suggests that both intracellular and extracellular Ca^2+^ contribute to processes extension, it indicates that the calcium-mediated cytoskeletal rearrangement is more likely initiated via membrane channels rather than from intracellular stores. A concentration ffect may also be taking place: a sustained influx from the medium may provide enough Ca^2+^ ion to elicit maximal morphological response, while a limited intracellular amount of calcium would only lead to a lesser extension. This implication makes the group I mGluRs, that induce intracellular calcium release from internal stores via the IP_3_ cascade, unlikely to be involved in processes extension caused by glutamate in U118-MG cells. This contrasts with the studies that have proposed mGluR5 (a group I mGluR) to be the main receptor in astrocytic glutamate signaling (Rose et al. 2018), and was found to be part of glutamate signaling in hippocampal astrocytes (Porter and McCarthy 1996; Latour et al. 2003; Panatier and Robitaille 2016). However, databases show the type of astrocytoma cell lines that we are using do not express GluR5 under normal cell culture conditions (Thul et al. 2017). In addition, our data acquired from U118-MG cells showing that extracellular calcium induces stronger filopodiagenesis, support previous studies that have questioned the function of astrocytic mGluR5 in glutamate signaling. For instance, mGluR5 agonists failed to induce calcium signaling in astrocytes in the adult brain (Sun et al. 2013). mGluR5 has also been reported to be only partially responsible for calcium signals in response to neural glutamate release in astrocytic processes from mature hippocampal mossy fibers in another research (Haustein et al. 2014). These studies combined to our observations suggest that a transmembrane calcium conductor must be involved within calcium events of astrocytic glutamate signaling (Oberheim et al. 2006; Verkhratsky and Kirchhoff 2007).

The membrane-bound ion channels iGluRs such as NMDA and AMPA receptors are obvious candidates for glutamate-induced filopodiagenesis. However, NMDA receptors have been indicated to be either unfunctional or absent in hippocampal cultured astrocytes or even in hippocampal slices (Matyash and Kettenmann 2010). In the type of astrocytoma cells that we are employing, the NMDA receptors shown to lack the essential NR1 subunit which makes them unfunctional (Swanson and Sakai 2009; Uhlen et al. 2010). On the other hand, calcium-permeable AMPA receptors have been reported to contribute to the calcium-dependent events associated with glutamate signaling: in Bergmann glial cells from cerebellar cortex (Matsui, Jahr, and Rubio 2005), in radial-like glial cells from the dentate gyrus subventricular zone (Swanson and Sakai 2009; Matyash and Kettenmann 2010; Rose et al. 2018), and in Bergmann glia appendages at Purkinje cell synapses (Swanson and Sakai 2009). However, U118-MG evidenced to express iGluR2 which restricts the Ca^2+^ conductance of AMPA receptors (Swanson and Sakai 2009; Uhlen et al. 2010) and therefore makes them unlikely to directly mediate the calcium influx required for cytoskeletal rearrangement following glutamate sensing.

Serendipitous observations in another part of our research program on astrocytic Cav’s led us to hypothesize that they have an active role in astrocytic glutamate signaling. Despite being non-electrical cells, astrocytes express voltage-gated calcium channels, however their function in these cells is still unclear (Latour et al. 2003; Cheli et al. 2016). We initially posited that astrocytic Cav’s might amplify the intracellular calcium amounts necessary for filopodiagenesis via actin reorganization. Upon glutamate stimulation, membranebound GluRs could activate Cav’_s_ by changing the membrane potential, therbery activating the voltage sensors of the pore-forming Cav’_s_ α1 subunit (Hille 2001; Youn, Gerber, and Sather 2013). Another possibility could involve adenylate cyclase-coupled mGluRs (mGluR8 in U118-MG (Thul et al. 2017)) that can potentiate Cav gating by phosphorylation (Hove-Madsen et al. 1996; Lepski et al. 2013).

The results from experiments with two sets of Cav inhibitors provide strong evidence for a functional role for astrocytic Cav’s in the morphological response of astrocytes to glutamate. Three inhibitors of Cav α1 significantly decreased the extent of glutamate-induced filopodia extension in U118-MG cells, but only partially (Fig. 2). In fact, a similar role for L-type Cav’s in the extension of neural dendrites has been reported recently (Kamijo et al. 2018; Zatkova et al. 2018). In these studies with neurons, the stimuli and downstream pathways differ from our study with astrocytoma cells, but the contribution of L-type Cav’s to the extension of processes is consistent. The limited filopodia extension observed with α1 inhibitors (nifedipine, verapamil, diltiazem) could be due to a parallel calcium-entry pathway sensitive to glutamate, or to secondary messenger signaling via intracellular calcium release. However, neither would appear to be sufficient to elicit a full morphological response.

Surprisingly, the morphological response of cells to glutamate was virtually abolished when gabapentinoids were ued, as inhibitors of α2δ1— Cav’s second subunit essential to calcium influx (Fig. 3). The extracellular α2δ subunit of Cav is known to modulate ion conductance of the α1 subunit, as well as to control its localization within the cell membrane (Woolley et al. 1990; Striessnig and Koschak 2008; Song et al. 2015; Robinson et al. 2011). In the context of neurons, α2δ has been confirmed to directly interact with the neuronal actin structures of the cell membrane (Robinson et al. 2010, 2011). It was also shown to affect the cytoskeletal rearrangement in neurons (Kurshan, Oztan, and Schwarz 2009), and to control dendritic spine morphogenesis (Christopher Risher et al. 2018). Thus, this Cav subunit is likely to act similarly in astrocytic morphology. Furthermore, α2δ has been demonstrated to be accumulated in brain where the excitatory glutamatergic synapses are formed. This specific localization may stregnthen a possible role in intercellular glutamate signaling events (Geisler et al. 2019). These studies converge with the results presented herein to suggest a significant role of α2δ protein in morphological responses of fine astrocytic processes.

It should be noted that gabapentinoids have been suggested to cause a Ca^2+^ influx in cells by enhancing sodium/glutamate co-transport of Na^+^/Ca^2+^ exchanger via its reverse mode (Kirischuk, Kettenmann, and Verkhratsky 1997; Rojas et al. 2007; Yoshizumi, Eisenach, and Hayashida 2012). If this side-effect of gabapentinoids occurred here, one would expect to see an increase of processes extension due to Ca^2+^ influx. Instead, gabapentinoids completely inhibited the morphological response of astrocytoma cells to glutamate. This unexpected strong inhibition suggests a different and currently undefined mechanism of action for gabapentinoids in abolishing glutamate-induced filopodiagenesis.

The study presented above opens a new avenue of research to decipher the contribution of astrocytic Cav subunits in functional synapses. It may also offer new insight into how gabapentinoid drugs act in the treatment of epilepsy and neuropathic pain (Taylor, Angelotti, and Fauman 2007; Robinson et al. 2011; Dolphin 2012; Chen et al. 2018). These findings are now leading us to further investigate the role of α2δ subunit of Cavs in PAPs motility and glutamate-dependent events within tripartite synapse.

## Acknowledgments

We are grateful to Andis Klegeris for generous gifts of cell lines.

## Authors contributions

MT conceived and performed the experiments, analyzed the data, and co-wrote the manuscript. FM conceived the idea and co-wrote the manuscript.

## COMPLIANCE WITH ETHICAL STANDARDS

### Ethical approval

This study did not involve protocols requiring ethical approval.

### Research involving human participants and/or animals

This Article does not contain any studies with human participants or animal performed by any of the authors.

### Informed consent

Informed consent was obtained from all individual participants included in the study.

